# fNIRS Characterization of Temporal Potentiation During Optimized Nerve Stimulation

**DOI:** 10.64898/2026.01.12.699125

**Authors:** Susan K. Coltman, William B. Riggs, Androu Abdalmalak, Xiaogang Hu

**Affiliations:** Department of Kinesiology, The Pennsylvania State University, University Park, PA, 16802 USA; Bioengineering Masters Program at the University of Pittsburgh, Pittsburgh, PA, 19122 USA; Independent Researcher; Department of Kinesiology, Department of Mechanical Engineering, Department of Physical Medicine and Rehabilitation, the Huck Institutes of the Life Sciences, and the Center for Neural Engineering, The Pennsylvania State University, University Park, PA, 16802 USA

**Keywords:** Functional near-infrared spectroscopy, Peripheral nerve, Sensory adaptation, Sensory feedback, Transcutaneous nerve stimulation

## Abstract

Restoring stable somatosensory feedback through transcutaneous nerve stimulation (TNS) represents a significant challenge in neural engineering; however, the cortical dynamics underlying sensory habituation are not yet well characterized. This study utilizes a multimethod functional near-infrared spectroscopy framework to quantify the temporal evolution of cortical engagement during optimized TNS. It was hypothesized that stimulation parameters fundamentally constrain the spatiotemporal profile of the hemodynamic response function. A General Linear Model with temporal and dispersion derivatives was used to analyze hemodynamic responses in 10 participants across three phases: validation, duration optimization (2, 5, or 10 seconds), and sustained engagement (45 minutes). During the optimization phase, 5-second stimulation produced the highest canonical amplitude with optimal dispersion characteristics, while 10-second trains resulted in significant temporal delays. During the sustained phase, parametric modeling indicated no linear decay in cortical response. Instead, the cortex demonstrated temporal potentiation, characterized by a transition from sparse early-phase activation to widespread late-phase recruitment across the sensorimotor network. The left posterior parietal cortex (BA5) served as a continuous integration hub, whereas primary somatosensory regions exhibited dynamic late-onset engagement. These findings quantify a nonlinear adaptation mechanism whereby engineered temporal spacing prevents neural saturation. The observed intersubject heterogeneity highlights the necessity for adaptive, closed-loop TNS controllers guided by real-time hemodynamic biomarkers.

## I. Introduction

TACTILE feedback is essential for daily activities such as grasping a coffee mug or operating a mobile phone. The loss of tactile input resulting from amputation, nerve injury, or neurological disorders impairs grip control and object manipulation, and fundamentally alters the brain’s representation of both the body and artificial limbs [1]–[4]. These deficits frequently contribute to prosthesis rejection and diminished quality of life [5], [6]. Transcutaneous nerve stimulation (TNS) is a promising, non-invasive approach to restore somatosensory input through electrical stimulation of median and ulnar nerve fibers [7], [8]. Evoked localized sensations can facilitate sensorimotor plasticity and improve neuroprosthetic performance [4], [9]. However, clinical translation depends on the ability to generate stable, meaningful sensations that persist over extended periods [10].

Prolonged or repetitive stimulation can cause rapid sensory habituation due to short-term adaptation in both peripheral and central pathways [11]–[13]. Our recent work demonstrated that optimizing the temporal structure of stimulation, specifically stimulation duration and rest intervals between stimuli, can substantially reduce perceptual habituation and sustain comfortable perception during extended stimulation [10]. While considerable progress has been made in understanding peripheral and behavioral responses, the central cortical mechanisms underlying sustained sensory encoding during TNS are still not well known. It is unclear how activity in the somatosensory and sensorimotor cortices changes with repeated stimulation or how timing affects these responses. To characterize TNS effects, we focused on somatosensory regions critical for tactile information processing: the primary somatosensory cortex (S1), secondary somatosensory cortex (S2), and posterior parietal cortex (PPC), which support touch perception and sensorimotor integration [14]–[16]. Understanding how these cortical regions respond to temporally structured stimulation is essential for developing reliable stimulation strategies.

Characterizing cortical adaptation during TNS requires methods that track neural activity over extended periods without restricting movement. Functional near-infrared spectroscopy (fNIRS) addresses this need by measuring brain hemodynamics via near-infrared light detection of oxygenated and deoxygenated hemoglobin [14], [17], [18]. fNIRS provides a non-invasive tool with a spatial resolution superior to electroencephalography (EEG) and temporal resolution superior to functional Magnetic Resonance Imaging (fMRI), making it well-suited for studying cortical activity during naturalistic behavior. [19], [20]. Although reconstructing precise cortical maps from optical measurements requires advanced signal-processing approaches [21], [47]–[50], cortical adaptation manifests as gradual shifts in hemodynamic amplitude that fNIRS can quantify non-invasively in somatosensory areas.

To address this gap, we designed a three-phase experiment that combines TNS and fNIRS to quantify how temporal parameters of stimulation modulate cortical activation across both short- and long-term timescales. Following cap validation (described in Methods), Block 2 examined TNS duration optimization, and Block 3 assessed sustained cortical engagement. This combined approach reveals how timing affects sustained sensory encoding in these regions [14], [22]–[24].

We hypothesized two competing but testable scenarios regarding cortical dynamics during prolonged TNS. First, we predicted that optimized temporal protocols would maintain relatively stable cortical activation throughout prolonged TNS. Early responses would reflect initial sensory encoding, while sustained late responses would indicate robust perceptual stability. Alternatively, we expected that spaced stimulation could trigger dynamic reorganization of cortical activation. In this case, the early-phase cortical activity would be minimal (reflecting initial adaptation), but delayed recruitment of higher-order sensorimotor areas would progressively engage over time. By examining both the magnitude and temporal evolution of hemodynamic responses across a sustained stimulation protocol, we aimed to distinguish between these mechanisms. Our goal was to establish temporal design principles for sustaining meaningful somatosensory engagement during sensory stimulation.

## II. Methods

### A. Participants

Ten neurologically intact adults (mean age of 26.8 years; age range 20-42 years; eight males) participated in this study. All the participants provided informed consent. This study was approved by the Institutional Review Board of the Pennsylvania State University (Approval Number: STUDY00022392).

### B. Electrical Stimulation Protocol

The upper right arm was cleaned with 70% isopropyl alcohol, abraded using an electrode preparation wipe to remove dead skin cells, and then wiped again with alcohol to ensure cleanliness prior to electrode placement. Following preparation, a 2×5 grid of 10 round electrodes (∼10 mm in diameter each) was placed on the medial side of the upper arm beneath the short head of the biceps brachii. The grid was positioned to target the median and ulnar nerves. Electrodes were arranged parallel to the underlying nerve pathways and spaced approximately 1.5 cm apart (**Fig. 1A**). Constant pressure on the electrode array was maintained by placing a foam pad, secured with surgical tape over the setup.

**Fig. 1.**
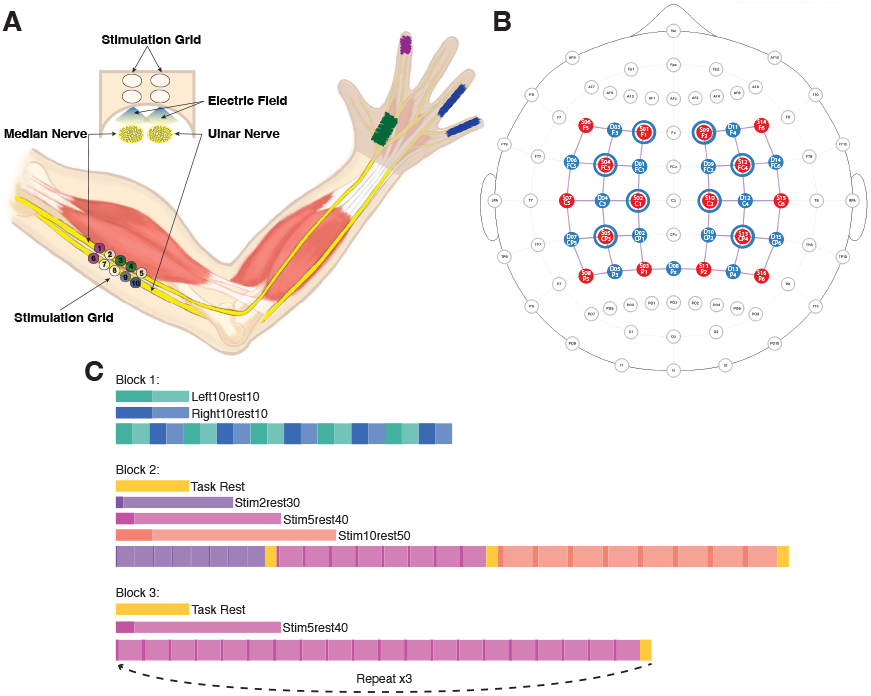
Experimental Setup and Protocol Schematic. (A) Peripheral Setup and Haptic Maps. This schematic illustrates the electrode grid placement on the left upper arm, targeting the median and ulnar nerves for transcutaneous nerve stimulation (TNS). The hand map presents specific palmar regions where perceived haptic sensation was evoked for three representative participants: P4, P7, and P10. (B) fNIRS Optode Montage. The 54-channel montage (16 Sources, 15 Detectors) targets the somatosensory and motor cortices, with 46 long-separation channels and eight short-separation channels; blue circles indicate the short-channel locations. (C) Temporal Block Design. The experimental protocol consists of two main blocks: Block 2 compares three TNS parameter conditions (stim2rest30, stim5rest40, stim10rest50) along with a task-rest baseline; Block 3 evaluates long-duration stimulation using the stim5rest40 trial structure, repeated 60 times. (Prior to these blocks, cap placement was validated using a finger-tapping localizer.)

After electrode placement, a 2-channel stimulator (STG4002, Multichannel Systems) was programmed to deliver charge-balanced square pulses, controlled by a switch matrix (Agilent Technologies) enabling dynamic electrode pair selection. A pulse frequency of 150 Hz and a pulse width of 200 *µ*s were chosen, based on studies supporting their effectiveness for stable, localized haptic sensations via TNS [25], [26]. These parameters remained constant to avoid confounding variability when assessing how temporal structures (e.g., stimulation duration, rest intervals) influence adaptation and habituation.

Prior to the main experiment, a systematic testing procedure was conducted using a custom MATLAB interface to identify optimal electrode pairs for selective nerve stimulation. Two functional pairs were identified for each participant that selectively evoked comfortable haptic sensations on the palmar side of the right hand, innervating either the ulnar or median nerve pathway while avoiding electrode discomfort or motor activation. Sensation thresholds were determined by incrementally increasing current from 1.5 mA until participants reported haptic sensation, then identifying motor threshold. Stimulation intensity for the main experiment was set at sensation threshold (range: 2.0-4.4 mA, Mean = 3.0 ± 0.7 mA) [26]. Electrode pairs elicited sensations in the median (e.g., Index, Middle, Thumb) or ulnar (e.g., Pinky, Ring) nerve distributions.

### C. fNIRS Signal Acquisition

fNIRS via NIRScoutXP (NIRx Medizintechnik GmbH, Berlin, Germany) was used to measure blood oxygen levels in the brain. The system utilized two wavelengths (760 and 850 nm) and acquired signals at a sampling frequency of 3.9 Hz. A montage was developed to determine the placement of the fNIRS sensors and decoders. The somatosensory cortex of the brain, the region where touch/cutaneous information is received, was specifically targeted [14]. We designed the probe layout to target the primary somatosensory cortex (S1; responsible for perception of touch), secondary somatosensory cortex (S2), primary motor cortex (M1; controls voluntary movement), and superior parietal lobule (BA5; area involved in sensory integration). A modified motor montage (16 sources × 15 detectors) was used, with the outermost row of optodes shifted posteriorly to extend coverage over the parietal regions. Eight short-separation channels (∼8 mm) captured superficial scalp hemodynamics for noise regression, resulting in a 54-channel setup: 46 long-distance cortical channels (∼30 mm) and 8 short-separation channels. The optode locations and targeted brain regions are shown in **Fig. 1B**.

After determining the two functional stimulation pairs, the fNIRS cap was placed on the participant’s head and calibrated. During the experimental procedure, participants were asked to remain still, focus on a fixation cross, and avoid focusing their thoughts on anything in particular to prevent data artifacts. Digital synchronization between stimulation delivery and fNIRS data acquisition was achieved using two protocols: the stimulation program communicated with the fNIRS computer using the lab streaming layer protocol, which, in turn, senttrigger pulses to the NIRx Scout recording system. This setup ensured accurate time-locking of all TNS onset times for accurate synchronization and subsequent General Linear Model (GLM) analysis.

Prior to the main experimental protocol, cap placement was functionally validated using a unilateral finger-tapping task. Participants performed five cycles of self-paced right-hand tapping (10 seconds of tapping, 10 seconds rest), followed by five cycles of left-hand tapping. Demonstration of clear contralateral activation in the primary motor cortex (BA4) was required to advance to the main experiment. This procedure confirmed that the fNIRS optode montage was accurately positioned to capture sensorimotor cortical activity.

### D. Experimental Design

#### 1) Pilot TNS Parameter Optimization (Block 2)

The second block served as an optimization task to select the TNS duration for the main, long-term protocol. The primary goal was to identify a pulse duration that reliably elicited a robust cortical hemodynamic response while minimizing early signs of habituation. Participants received three separate sub-blocks of TNS delivered in a pseudo-randomized order, with each sub-block employing a different stimulation/rest timing ratio. The conditions included Short TNS (2s stimulation with 30s rest; stim2rest30), Medium TNS (5s stimulation with 40s rest; stim5rest40), and Long TNS (10s stimulation with 50s rest; stim10rest50). Each ratio was repeated eight times within its sub-block. Throughout all stimulation and rest periods, the participants were instructed to remain still and maintain a resting state focus. A 20s rest break was provided between the three sub-blocks to allow for brief adjustments or communication, which was followed by a 5-minute break before proceeding to the main experiment (see **Fig. 1C** for a schematic representation of the timing ratios).

#### 2) Main Long-Duration TNS Protocol (Block 3)

The main block used a 5-second TNS duration (5s stim, 40s rest; stim5rest40) based on pilot data identifying this as optimal for a strong hemodynamic response. This condition repeated 60 times over 45 minutes, split into three 20-rep segments with short breaks. Participants stayed still and maintained resting focus throughout.

### E. Data Analysis and Statistical Analysis

The fNIRS data were preprocessed and analyzed using the NIRS Brain AnalyzIR Toolbox[26] implemented in MATLAB R2024a. The analysis pipeline consisted of individual-level (first-level) and group-level (second-level) GLM.

#### 1) Data Quality Control

Signal quality was assessed using a two-tiered coefficient of variation (CV) threshold applied to the raw light intensity data. First, an exclusion threshold (CV *>*7.5%) was applied across the entire run to flag low-quality channels, following established guidelines [28], [29]. Second, individual trials were excluded if *>*10% of the channels were contaminated. Across the remaining 10 participants, 18 of 1,208 trials (1.49%) were excluded, indicating an overall high signal quality.

#### 2) Preprocessing

Raw light intensity data were first converted to optical density, and motion artifacts were corrected using Temporal Derivative Distribution Repair (TDDR; [30]). Signals were then converted to oxyhemoglobin (HbO) and deoxyhemoglobin (HbR) concentration changes using the modified Beer–Lambert Law [31]. To suppress slow drifts and high-frequency noise, a band-pass filter (0.01–0.5 Hz) was applied using the HOMER2 hmrBandpassFilt function [20], [32].

#### 3) First-Level GLM

For each participant, a GLM was fitted to estimate the task-evoked hemodynamic response at each channel. All models employed robust regression using an autoregressive iteratively reweighted least squares (AR-IRLS) model to reduce the impact of outliers and serial correlations [20], [27]. Short-separation regressors were included to control for superficial hemodynamics through implementation of nearest short-separation channel inclusion within the GLM, and third-order Legendre polynomials were used to model the slow temporal trends [27], [33], [34]. To accurately model the temporal dynamics of the hemodynamic response, we employed a GLM with temporal and dispersion derivatives. This approach aligns with established signal modeling frameworks in medical imaging [35] that account for the sluggish nature of the hemodynamic response [36]. In addition, BoxCar basis functions were used to model the 20-second task rest periods between stimulation series, allowing task-evoked activity to be isolated from unstructured variance, such as movement or baseline drift [37]. The specific basis functions, regressors of interest, and subsequent *β*-estimates, which quantify the magnitude and temporal characteristics of the evoked hemodynamic change, varied across blocks and analytical modes.

#### 4) Second-Level Group Analyses

To determine group-level effects, individual-level *β*-estimates were entered into a mixed-effects analysis of variance (ANOVA) using the AnalyzIR Toolbox [27]. This model incorporated fixed effects of condition, random intercepts for each participant, and full channel-level beta covariance matrices to account for serial correlations and between-participant variability. False discovery rate (FDR) correction (*q <* 0.05) was applied to all channels.

#### 5) Region-of-Interest (ROI) Analysis

To complement the channel-level analysis, an ROI analysis was performed. Group-level *β*-estimates were averaged within anatomically defined ROIs specific to the tasks performed in each block. For Blocks 2 and 3, the analysis included the left and right primary somatosensory (BA2, BA3), secondary somatosensory (BA40, BA43), and posterior parietal (BA5) regions, reflecting the somatosensory nature of the stimulation [14]. Statistical significance at the ROI level was assessed using the same mixed-effects GLM framework employed in the group-level analysis, with an FDR correction applied to control for multiple comparisons.

#### 6) Visualization

Source and detector locations were coregistered to the Colin27 head model in AtlasViewer [38], [39], and probe sensitivity profiles were estimated using Monte Carlo light transport simulations. Group-level activation maps were rendered onto a standardized Colin27 cortical mesh using image reconstruction with a Gaussian spatial basis (20 mm FWHM) [40]. Three-dimensional projections were generated to visualize the activation of the topography.

## III. Results

Functional validation of the fNIRS cap placement using the finger-tapping task confirmed the anticipated contralateral sensorimotor activation, thereby supporting the validity of the optode montage. The following results focus on cortical responses to transcutaneous nerve stimulation during the optimization (Block 2) and sustained engagement (Block 3) phases. Block 2 examined how TNS duration (2s, 5s, and 10s) modulates cortical hemodynamic responses. We evaluated the effects of stimulation duration on the amplitude, timing, and spatial distribution of HbO activation to identify optimal parameters for the main sustained stimulation protocol.

### A. Canonical Hemodynamic Response Function (HRF) Amplitude (β01)

The canonical HRF beta (*β*01) primarily represents the overall magnitude of the hemodynamic response. Consistent with our first hypothesis, all stimulation durations produced significant increases in HbO concentration across the somatosensory and parietal regions (all *q* <0.05), confirming that even brief TNS trains reliably engaged cortical areas associated with tactile processing. The strongest canonical amplitude activations for each condition are presented in **Table I** The maximal canonical response overall was elicited by the 5s TNS at channel S1-D3 (F(1,8025) = 24.96, *q* = 0.0002). The 2s TNS and 10s TNS conditions produced their highest canonical effects in S3-D8 and S1-D3, respectively, with full details provided in **Table I**.

**TABLE I.** Peak Statistical Channels BY Condition AND GLM Component.

| Condition | $\beta$ Component | Description | Chan. | F-stat | q-val |
| --- | --- | --- | --- | --- | --- |
| stim2rest30 | Canonical ($\beta_{01}$) | Response Amplitude | S3-D8 | 19.317 | 0.00206 |
| stim5rest40 | Canonical ($\beta_{01}$) | Response Amplitude | S1-D3 | 24.959 | 0.00022 |
| stim10rest50 | Canonical ($\beta_{01}$) | Response Amplitude | S1-D3 | 14.483 | 0.00752 |
| stim2rest30 | Temporal ($\beta_{02}$) | Timing Shift (Delay) | S11-D8 | 18.243 | 0.00302 |
| stim5rest40 | Temporal ($\beta_{02}$) | Timing Shift | S13-D15 | 12.514 | 0.01483 |
| stim5rest40 | Dispersion ($\beta_{03}$) | Response Shape (Prolonged) | S10-D12 | 17.589 | 0.00364 |

The table highlights the single most statistically significant channel for each GLM component across the three TNS duration conditions (2s, 5s, and 10s). The canonical component ( $\beta_{01}$ ) estimates the overall response amplitude. The temporal component ( $\beta_{02}$ ) captures shifts in response timing. The dispersion component ( $\beta_{03}$ ) captures changes in the response shape or duration.

To clarify the differences in amplitude between conditions, we directly contrasted the canonical HbO betas (*β*01) for each TNS duration, with FDR correction for multiple comparisons (*q* < 0.05). No significant difference was found in HbO amplitude between 2s TNS and 5s TNS in any channel, indicating both durations produced similar maximum cortical responses. In contrast, the comparison of 10s TNS versus 5s TNS revealed several channels where HbO amplitude was significantly greater for the 10s TNS condition. Specifically, in the somatosensory channels S3-D5 (Contrast *β* = 7.78, *q* = 0.012) and S11-D8 (Contrast *β* = 7.09, *q* = 0.0165), as well as the parietal site S11-D13 (Contrast *β* = 7.66, q = 0.012), 10s TNS produced a larger amplitude than 5s TNS. These results indicate that while 2s and 5s TNS reach similar peak amplitudes, extending the stimulation to 10s drives a higher magnitude of cortical activity in specific somatosensory and parietal regions.

### B. Temporal (β02) and Dispersion (β03) Derivatives

In addition to overall activation amplitude, analysis of derivative terms demonstrated that stimulation duration modulates both the timing and shape of the cortical response. The temporal derivative (*β*02) quantifies the latency of the response peak; positive values indicate a delay relative to the canonical model. Significant *β*02 effects were observed across all stimulation durations, indicating a duration-dependent shift in response timing. As shown in **Table II**, the most prominent shifts were localized to S11-D8 for 2s TNS (*F* = 18.24, *q* = 0.003), S13-D15 for 5s TNS (*F* = 12.51, *q* = 0.015), and S4-D4 for 10s TNS (*F* = 14.86, *q* = 0.007). These findings demonstrate a systematic trend: increasing stimulation duration results in a delayed, temporally extended hemodynamic response. This pattern is consistent with the fMRI literature, in which longer tactile stimuli elicit later and more prolonged HbO response peaks. Accurately characterizing these timing features is essential for engineering applications such as closed-loop neuromodulation, which require precise temporal alignment with the user’s brain state.

**TABLE II.** Key Statistically Significant Findings FROM Group-Level ROI Analysis.

| Cond. | $\beta$ Comp. | ROI | $\beta$ Est. | q-val | Interpretation |
| --- | --- | --- | --- | --- | --- |
| stim5rest40 | Canonical ($\beta_{01}$) | BA2_left (S1) | 1.313 | 0.0036 | Strongest overall activation. |
| stim2rest30 | Temporal ($\beta_{02}$) | BA43_right (S2) | -5.325 | 0.0208 | Indicates significant timing shift for short TNS. |
| stim5rest40 | Temporal ($\beta_{02}$) | BA40_right (S2) | -2.561 | 0.0021 | Indicates significant timing shift for moderate TNS. |
| stim5rest40 | Dispersion ($\beta_{03}$) | BA40_right (S2) | 2.293 | 0.0349 | Confirms sustained S2 response for 5s TNS. |
| stim5rest40 | Dispersion ($\beta_{03}$) | BA3_right (S1) | 2.263 | 0.0438 | Confirms sustained S1 response for 5s TNS. |

Cond. = Condition; Est. = Estimate. Values demonstrate duration-dependent effects across primary (S1) and secondary (S2) somatosensory cortices.

The dispersion derivative (*β*03) reflects the duration or width of the hemodynamic response, indicating how long the activity is sustained. In single-channel analyses, significant dispersion effects were observed only for the 5s TNS condition, primarily due to a strong effect at channel S10-D12 (*F* = 17.59, *q* = 0.004). However, this channel-specific perspective may overlook broader patterns. When region-of-interest (ROI) level aggregation was employed to enhance statistical power, a more extensive pattern of modulation was detected. Significant dispersion effects were found in multiple regions, including the right primary (BA3) and secondary (BA40, BA43) somatosensory cortices for both the 2s and 5s TNS conditions. These results support the hypothesis that TNS duration systematically modulates the sustained characteristics of the cortical response, even when these effects are spatially distributed rather than localized to a single channel.

### C. Spatial Modulation of Response

Finally, we examined whether different stimulation durations recruited distinct regions of the cortex. **Fig. 2** illustrates that the spatial distribution of significant channels (and their corresponding HRF component) differed between durations: 2s stimulations primarily activated midline and contralateral somatosensory regions (e.g., S3-D8, S10-D12); 5s stimulations produced more widespread bilateral activation, including posterior parietal sites; and 10s stimulations engaged a subset of overlapping but distinct channels within the S1 and parietal cortex. This pattern suggests that stimulation duration modulates not only the magnitude and timing but also the spatial organization of cortical recruitment. Collectively, the model-based analyses revealed clear duration-dependent effects in timing, shape, and spatial distribution that were obscured in simple averaged responses, highlighting that TNS duration could modulate when, how long, and where cortical activation unfolds.

**Fig. 2.**
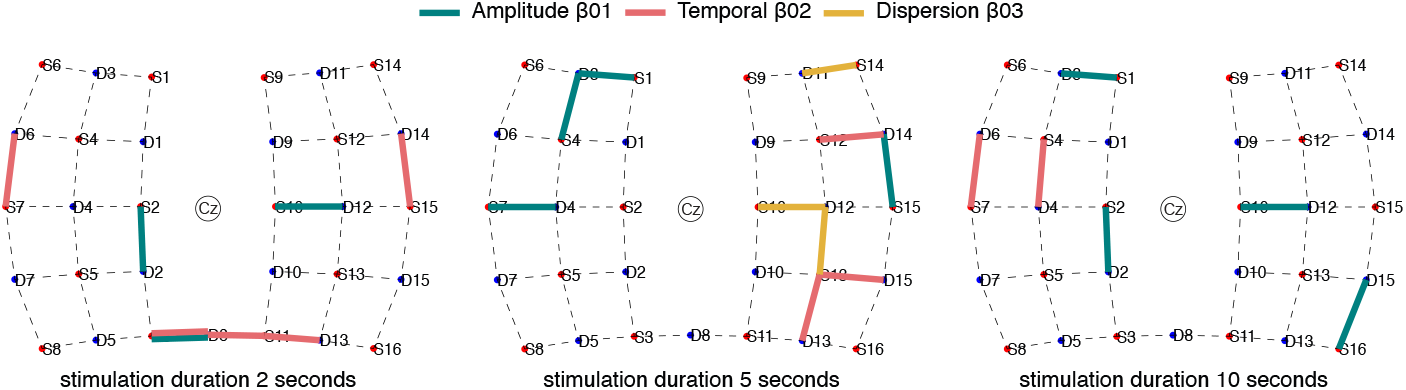
Duration-Dependent Spatial Modulation of HRF Components. Illustrating all channels that exhibited statistically significant HbO changes (q ***<*** 0.05 FDR-corrected) for the 2s, 5s, and 10s TNS durations. Significant effects are color-coded by the GLM component: Amplitude (***β***01, teal), temporal derivative (***β***02, coral), and dispersion derivative (***β***03, mustard).)

### D. ROI Group Analysis

To complement these channel-level findings and confirm that duration-dependent effects generalize across functionally defined brain regions, we performed an ROI-based analysis aggregating responses within anatomically defined somatosensory and parietal regions. All selected ROIs showed significant canonical (*β*01) HbO activation for at least one TNS duration. This confirms that the canonical amplitude effect observed at the channel level is robust and generalized across the entire somatosensory-parietal network being studied. The strongest overall canonical response was observed in the left primary somatosensory cortex (BA2 left) for the 5s TNS condition (*β* = 1.313, *q* = 0.0036; **Table II**).

The dispersion effect (*β*03) was widespread. While previously confined to a single channel (S10-D12), regional analysis found significant dispersion effects in multiple ROIs, including BA3 right, BA40 right, BA43 right, BA43 left, and BA5 left. This effect was significant for both the 2s and 5s TNS conditions. As shown in **Table II**, the 5s TNS condition produced significant dispersion effects in both primary (BA3 right, *β* = 2.263, *q* = 0.0438) and secondary (BA40 right, *β* = 2.293, *q* = 0.0349) somatosensory cortices. This is a major finding that validates the hypothesis that TNS duration systematically modulates the sustained nature of cortical hemodynamic response.

Finally, temporal shifts (*β*02), which indicate changes in the timing of the response peak, were widespread. Significant temporal derivative effects were observed across almost every ROI for at least one condition, supporting the conclusion that the timing of the hemodynamic response was duration-dependent. Notably, the 2s TNS produced the largest timing shift in the secondary somatosensory cortex (BA43 right), while the 5s TNS showed more moderate shifts (**Table II**). Overall, the ROI analysis provides strong evidence that TNS duration does not merely affect a single point in the brain but induces systematic and differentiable temporal and spatial modulations across a broad cortical network.

Based on these optimization results, particularly the robust canonical response and unique dispersion characteristics of the 5s condition, we selected this stimulus condition for the main experimental protocol (Block 3). This extended 45-minute session allowed us to investigate whether repeated TNS exposure would lead to habituation (decreased response over time) or sustained cortical engagement, a critical consideration for TNS systems designed for continuous or long-duration operations, such as prosthetic sensory feedback or therapeutic neuromodulation.

### E. Sustained Cortical Activation and Temporal Dynamics

The main effect of 5-second stimulation maintained a strong activation over the entire 45-minute period. The canonical component (*β*01) revealed highly significant increases in HbO concentration across 18 somatosensory and parietal channels (*q <* 0.05 FDR-corrected, F(1,5619) up to 70.73), with the most pronounced effect at channel S2-D2 (F(1,5619) = 70.73, *q <* 0.001). The temporal derivative (*β*02) was significant in eight channels, confirming that even with sustained stimulation, the response timing remained distinct from the canonical HRF. The dispersion derivative (*β*03) showed a significant effect in a single channel. This indicates that while the HRF shape (width/duration) was consistent with the canonical model across most channels, one localized region exhibited a statistically distinct average shape.

Contrary to the hypothesis of linear temporal evolution, the parametric term for the canonical amplitude (*β*01) was not significant in any channel. This finding suggests that the overall magnitude of the cortical response did not decrease linearly over the 45-minute period. While the amplitude remained stable, a highly localized adaptation was observed in response timing. The parametric term for the temporal derivative (*β*02) was significant at a single somatosensory channel, S3-D8 (F(1,5619) = 11.40, *q* = 0.0111). This suggests that although the response intensity was maintained, the peak timing of the hemodynamic response in this localized region shifted significantly and linearly (delay or acceleration) over a long duration. Finally, the parametric term for the dispersion derivative (*β*03) also yielded no significant channels, confirming that the sustained nature (width/duration) of the individual HRF did not exhibit a linear change across the 45-minute block. The absence of progressive amplitude decline indicates that TNS can maintain consistent cortical activation over extended durations, supporting its feasibility for applications requiring sustained sensory feedback.

### F. ROI Analysis Summary

Aggregating the channel results by Brodmann Area confirmed the widespread, highly significant canonical amplitude (*β*01)across the somatosensory and parietal cortices. As summarized in **Table III**., the strongest overall response was found in the Left BA5 (F(1,5619) = 70.73, *q* < 0.0001). Significant canonical effects were also observed bilaterally in BA2 and BA3, as well as in the Right BA43. Furthermore, significant temporal derivative (*β*02) effects were found in the Left BA5 (F(1,5619) = 17.19, *q* = 0.0005) and Left BA3 (F(1,5619) = 14.96, *q* = 0.0013), indicating variation in HRF timing across these regions. The parametric regressor designed to capture a linear decrease in amplitude (temporal evolution term) was not significant across any of the aggregated Brodmann Areas.

**TABLE III.** Summary OF Strongest Canonical Amplitude Effects IN Sustained TNS.

| Brodman Area | Hemi. | Component | F(1,5619) | q-val |
| --- | --- | --- | --- | --- |
| BA5<br>(Superior Parietal Lobule) | Left | Canonical Amplitude | 70.73 | < .0001 |
| BA43<br>(Secondary SC) | Right | Canonical Amplitude | 20.53 | .0001 |
| BA2<br>(Primary SC) | Left | Canonical Amplitude | 16.31 | .0008 |
| BA3<br>(Primary SC) | Left | Canonical Amplitude | 14.99 | .0013 |

Hemi. = Hemisphere; SC = Somatosensory Cortex. Values represent the strongest significant HbO effects during the sustained TNS phase.

### G. Temporal Analysis of Trial Bins

While the linear model suggests overall stability, a finer-grained analysis was required to detect potential non-linear temporal dynamics. Binning the data over time segments revealed a clear nonlinear temporal modulation in the cortical HRF. This pattern was characterized by minimal activity in the early phase, followed by a substantial increase in both the spatial extent and statistical significance of activation in the later bins. Specifically, the initial cortical engagement was sparse and highly localized across the first 20 trials (**Fig. 3**). In the Early Phase (bin 1, trials 1-10, and bin 2, trials 11-20), only a single channel at S2-D2 showed a significant increase in HbO (*q <* 0.05). The strongest statistical effect was observed in bin 1 at S2-D2, with a highly significant F-value (F(1,5535) = 18.574, *q* = 0.0021). This channel remained the sole significant response in bin 2, albeit with a slightly lower F-value (F(1,5535) = 13.526, *q* = 0.0127).

**Fig. 3.**
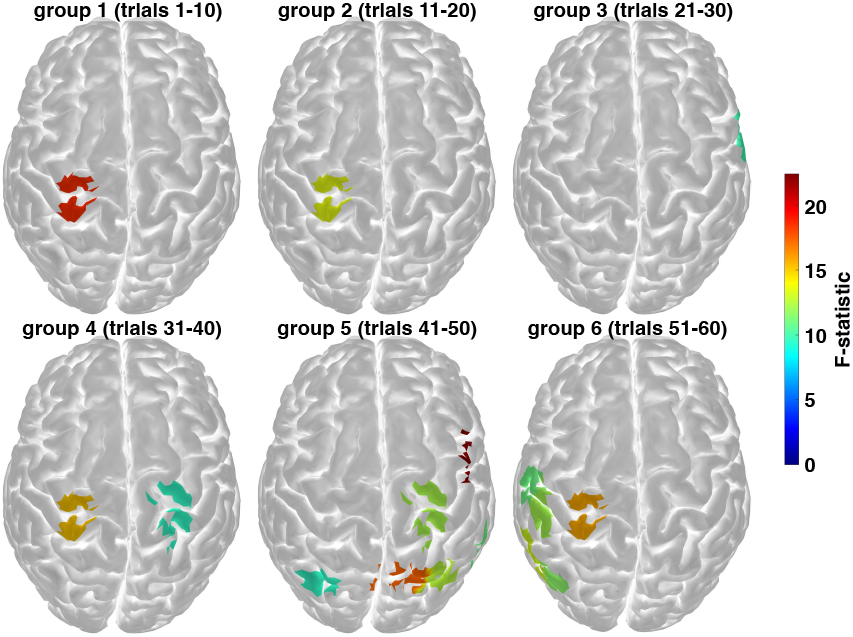
Temporal evolution of F-statistics for HbO activation across six temporal bins. This figure displays the activation maps (top-down view) for long-distance fNIRS channels across the motor cortex, grouped by bins of trials (bin 1-6). The data represent F-statistics derived from the GLM analysis for HbO changes during the task. The color scale, shown on the right, indicates the magnitude of the F-statistic, ranging from 0 (non-significant/below threshold) to the maximum observed value. Activation is only displayed for channels where the FDR corrected *q* < 0.05.

Activation remained minimal through Mid-Session Fluctuation. Bin 3 (trials 21-30) showed only a single significant channel (S15-D14, F(1,5535) = 9.9717, q = 0.0332). Bin 4 (trials 31-40) saw a slight re-engagement of the S2-D2 signal (F(1,5535) = 15.205, q = 0.0063) alongside one additional significant channel (S10-D10, F(1,5535) = 9.4969, q = 0.0391), totaling two significant sites.

A marked and widespread potentiation was observed in the late experimental session (bins 05 and 06), representing the final third of the protocol. Group 05 (trials 41-50) demonstrated the highest level of cortical recruitment, with six distinct channels showing a significant increase in HbO. The most robust effect in this group was found at S12-D12 (F(1,5535) = 22.413, q = 0.0007). This widespread response was sustained through group 06 (trials 51-60), which maintained five significantly activated channels. The strongest effect in the final group was observed at S2-D2 (F(1,5535) = 15.798, q = 0.0051). This finding suggests that the TNS-induced cortical response exhibits delayed recruitment or temporal potentiation over the course of the session rather than linear adaptation or fatigue.

### H. ROI-Based Temporal Analysis

The analysis confirmed the overall pattern of temporal potentiation observed in the channel-wise data and further highlighted the specific cortical regions driving this effect. The results revealed that the Left Posterior Parietal Cortex (BA5_left) was the most robust and consistently activated region across the entire 45-minute TNS session. Unlike the sporadic channel-wise activity, BA5 left showed significant activation in every group from group 01 to group 06 (**Fig. 4A**). The most dominant statistical effect of the entire session was observed in the first group (group 01) within this region, showing a highly significant HbR decrease (F(1,5535) = 21.782, *q* = 0.0004) and an HbO increase (F(1,5535) = 18.574, *q* = 0.0012). This sustained activation pattern in BA5 left suggests that this region is critical for processing tactile input throughout the entire duration. Despite this robust group-level response, individual subject trajectories demonstrated a certain level of variability across temporal bins as expected (**Fig. 4C**).

**Fig. 4.**
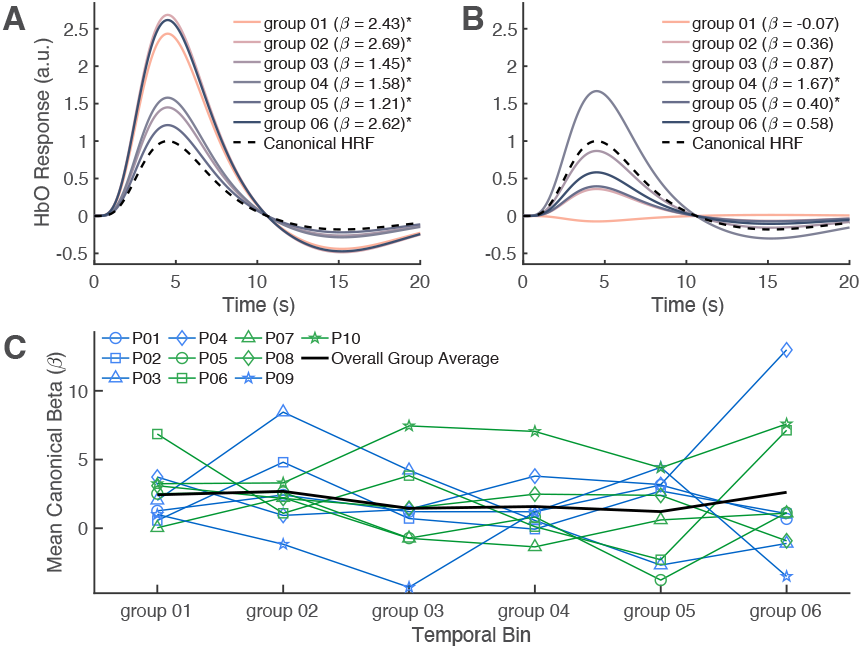
Hemispheric Comparison of Temporal Activation in PPC (BA5). A: Modeled canonical HRF amplitude (***β*** values) for the left BA5, plotted across six temporally binned groups of the 5-second TNS protocol. B: Corresponding ***β*** values for the right BA5, shown using the same temporal bins for direct comparison. C: Individual subject trajectories plotted across temporal bins with overall group mean (black line). Blue lines/symbols indicate ulnar nerve stimulation; green indicate median nerve stimulation. Significant effects are indicated with an asterisk (*).

Other somatosensory regions showed sporadic and later recruitment patterns. The right hemisphere, specifically BA43 right (S2), showed a strong initial HbR decrease in group 01 (F(1,5535) = 17.262, *q* = 0.0015) but failed to sustain a significant effect. The strongest recruitment outside BA5 left occurred in the Late Phase, with regions such as BA3 right (S1) showing significant HbO activation in group 05 (F(1,5535) = 9.193, *q* = 0.0263), and BA40 left (S2) showing significant HbO activation in group 06 (F(1,5535) = 7.461, *q* = 0.0402).

### I. Somatotopic Specificity and Temporal Binning (Median vs. Ulnar)

As an exploratory analysis to assess the specificity of the temporal potentiation pattern, we separated the data by nerve type (median n=5; ulnar n=5). Median nerve stimulation revealed a characteristic temporal potentiation pattern (**Fig. 5A**). Early activation was highly localized (group 01: single channel at S2-D2, F(1,2471) = 12.431, *q* = 0.0198) with intermittent mid-session activity (groups 02-04:0-2 significant channels). The late phase (groups 05-06) demonstrated marked increases in cortical recruitment, with group 05 showing six distinct significant channels (peak at S11-D8, F(1,2471) = 22.373, *q* = 0.0008), and group 06 showing four channels. ROI analysis confirmed this pattern, with left BA5 showing significant activation from group 01 onwards, expanding bilaterally in group 04, and recruiting left BA40 and BA2 in group 06.

**Fig. 5.**
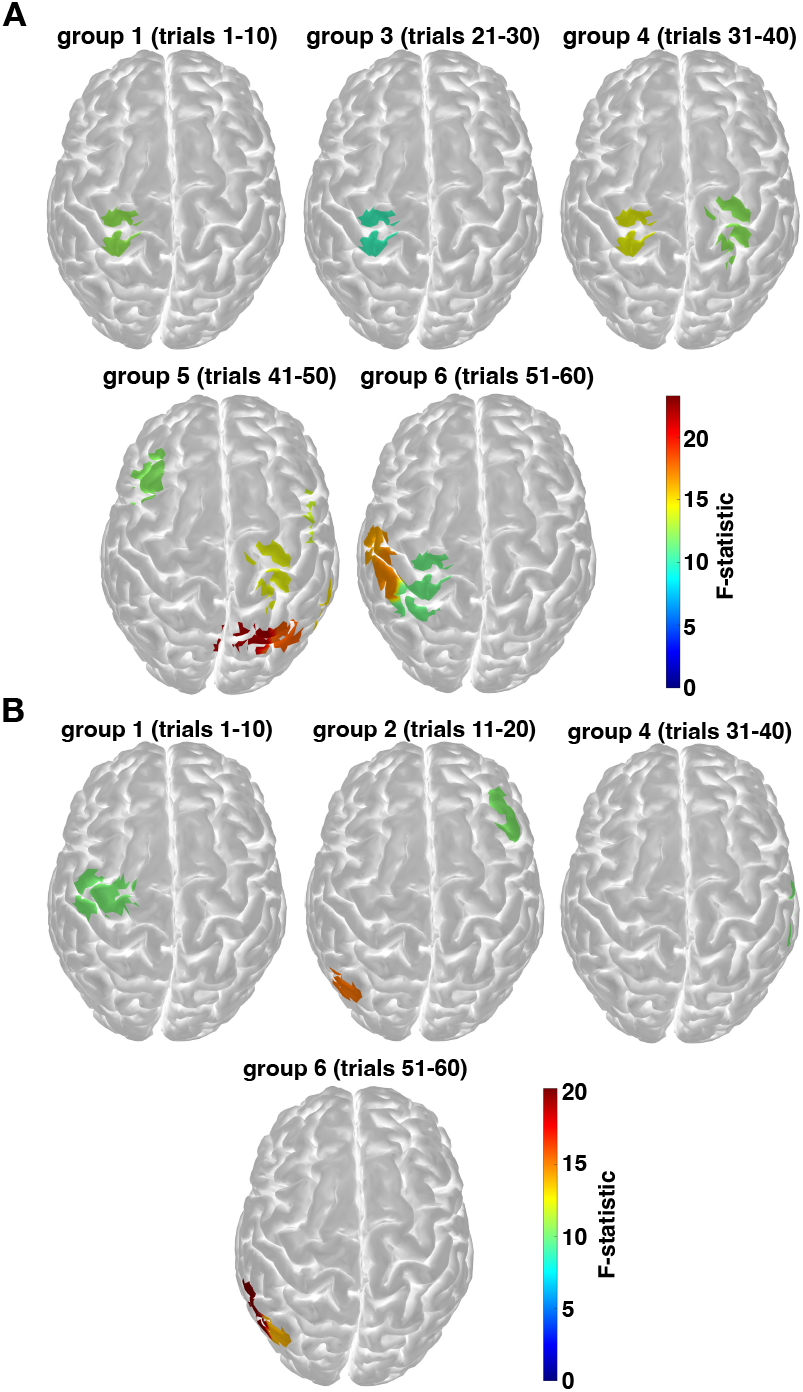
Temporal evolution of F-statistics for HbO activation during A: median and B: ulnar nerve stimulation. Surface projection maps (top-down view) illustrate the time course of HbO activation in the motor cortex as measured by long-distance fNIRS channels during the respective stimulation tasks. Significant channels (FDR-corrected *q* < 0.05) are colored, while non-significant channels show the underlying brain surface.

In contrast, ulnar nerve stimulation produced more sparse activation (**Fig. 5B**). Channel-wise analysis revealed intermittent significant effects across groups (group 01: one channel; group 02: two channels; groups 03-05:0-1 channels; group 06: two channels, peak at S8-D7, F(1,2428) = 20.257, q = 0.003). No ROI-level effects reached significance for any time bin. These nerve-specific differences are relevant for selective nerve-targeting strategies in TNS-based sensory systems, although the presence of late-phase activation in both pathways supports the primary finding of temporal potentiation rather than habituation.

## IV. Discussion

We implemented a multi-dimensional fNIRS approach to characterize the cortical dynamics of TNS across varying temporal scales. During the optimization phase (Block 2), stimulation duration was found to influence not only the magnitude but also the timing and spatial organization of the hemodynamic responses. All tested durations activated the somatosensory and parietal cortices; however, the 5-second duration achieved the optimal balance, producing the strongest canonical amplitude in S1 and distinct dispersion characteristics in sensorimotor regions. In contrast, 10-second stimulations resulted in significant temporal delays and prolonged dispersion. These findings align with foundational work establishing that stimulation parameters fundamentally constrain the spatial distribution [16] and temporal dynamics [41] of cortical activation.

During the sustained stimulation phase (Block 3), the optimized temporal protocol maintained cortical engagement as expected. Parametric analysis indicated no significant linear decline in amplitude across the 45-minute session, confirming that the selected rest intervals effectively prevented habituation. Importantly, the cortex did not remain static but exhibited temporal potentiation characterized by dynamic reorganization. Sparse, localized activity in the early phase (trials 1 to 20) progressed to widespread, robust recruitment of the sensorimotor network in the late phase (trials 41 to 60). These findings indicate that spaced stimulation facilitates delayed recruitment of higher-order areas rather than merely preventing decay.

Although robust group-level potentiation was observed, especially in the left posterior parietal cortex (BA5), individual subject trajectories exhibited considerable heterogeneity. As shown in **Fig. 4C**, there was no consistent ordinal ranking of responders across temporal bins, and this variability was observed across both nerve types (median versus ulnar). These results indicate that, while late-stage potentiation is a general phenomenon, the temporal profile of adaptation is highly individualized and likely involves mechanisms beyond peripheral nerve recruitment [23].

### A. Interpretation and Mechanistic Insights

The lack of a negative linear trend in canonical amplitude (*β*01) effectively rules out simple fatigue or habituation as the primary mechanism underlying cortical dynamics during optimized TNS. However, the presence of localized timing shifts indicates that, although the overall response magnitude is preserved, specific sensorimotor circuits are subject to fine-tuning in their timing and engagement patterns. The shift from minimal early engagement to wider recruitment suggests that early TNS inputs might be filtered, but longer, spaced-out exposure activates higher-order sensorimotor circuitry. Notably, we observed bilateral activation in S2 and PPC even under unilateral stimulation, consistent with known somatosensory processing networks [51]. Recent behavioral findings have quantified the rapid decay of sensory perception during continuous nerve stimulation [25], [46]. Consistent with these observations, new evidence suggests that optimized temporal spacing permits neural circuitry to re-engage, thereby maintaining perceptual stability [10].

The absence of a linear amplitude decline over the 45-minute session holds particular significance for clinical translation. These results demonstrate that temporally optimized TNS can maintain cortical engagement at clinically relevant levels, which is essential for continuous prosthetic use over extended periods. The observed hierarchical engagement pattern, with early-phase filtering followed by late-phase widespread recruitment, offers a mechanistic explanation for the failure of continuous stimulation due to neural saturation [11]–[13], [42], while highlighting the efficacy of spaced stimulation in facilitating repeated hierarchical re-engagement. These findings directly validate previous temporal optimization studies and provide a principled rationale for TNS system design.

Importantly, the temporal potentiation pattern was observed for both median and ulnar nerve stimulation, although the median nerve exhibited more robust region-of-interest effects. The generalization of temporal potentiation across both major hand sensory pathways suggests that this is a fundamental property of TNS-evoked cortical dynamics, rather than a pathway-specific phenomenon. This finding enhances the applicability of the identified design principles to a range of peripheral targets in sensory neuroprosthetics [1], [9].

A key finding is the regional specificity of the left posterior parietal cortex, which consistently functioned as the anchor of cortical activation throughout the session. In contrast, both primary and secondary somatosensory cortices demonstrated only sporadic or late-phase recruitment. This dissociation indicates a hierarchical temporal architecture. The posterior parietal cortex engages continuously to integrate artificial sensory input, which may establish a stable body schema representation [3], [23], [24], [43] distributed sensory processing areas are recruited dynamically as the session progresses. The robust group-level response in posterior parietal cortex, despite heterogeneous individual trajectories, suggests the presence of a stable underlying mechanism suitable for clinical deployment. This mechanism also offers potential for personalization to optimize individual outcomes [18].

The divergence in individual trajectories, despite the presence of a general mechanism, raises the question of why individuals adapt differently. Potential contributing factors include variations in peripheral sensitivity (observed here with sensation thresholds ranging from 2.0 to 4.4 mA), differences in central adaptation rates, or fluctuations in attentional state [12], [20], [25], [26]. While the within-subject repeated-measures design offers substantial statistical power for characterizing temporal dynamics, the sample size (n=10) limits the ability to draw definitive mechanistic conclusions.

### B. Clinical Implications

These findings establish essential design principles for sensory neuroprosthetics. The Block 2 optimization demonstrates that a 5-second stimulation window with sufficient rest (40 seconds) preserves signal integrity and avoids the temporal delays associated with longer stimulation trains. This delay is consistent with fMRI findings, where prolonged tactile stimuli result in shifted hemodynamic peaks [41]. For clinical translation, these results indicate that continuous stimulation strategies are likely suboptimal; temporal spacing of stimulation is necessary to enable cortical circuits to reset and engage hierarchically rather than undergoing habituation [10]–[13], [42].

Additionally, the inter-subject heterogeneity challenges the effectiveness of a universal approach to TNS parameterization. The observation that users do not respond uniformly to the same optimized protocol indicates that sensory stimulation protocols should be personalized based on individual cortical dynamics rather than solely on nerve anatomy [1], [9]. The variability documented in Figure 4C highlights the necessity for personalized, adaptive strategies. The loss of consistent tactile input is a recognized factor in prosthesis rejection and reduced quality of life [44]. These findings support the development of closed-loop adaptive systems [17], [18], [45], in which real-time monitoring of hemodynamic biomarkers [39], [40], such as BA5 recruitment, could dynamically adjust stimulation parameters to maintain cortical engagement [43].

### C. Limitations

Several limitations necessitate caution in generalizing these results. Although the sample size (n=10) is modest for population-level inferences, the within-subject repeated-measures design, with 60 trials per participant divided into six temporal bins, provides substantial statistical power to characterize both individual- and group-level temporal dynamics [27], [32]. The exploratory nerve-type analysis (n=5 per group) is limited and requires larger cohorts for definitive conclusions. Furthermore, fNIRS is restricted to cortical surface imaging, which precludes assessment of deep sulcal or subcortical contributions, such as thalamic gating, to the observed potentiation [20], [32]. The single-session design also leaves unresolved whether temporal potentiation persists or consolidates across multiple days of stimulation. Finally, although strict sensation thresholds were maintained, behavioral perceptual ratings were not collected during the long-duration block, limiting the ability to directly correlate cortical potentiation with sustained perceptual intensity.

### D. Future Directions

Future research should prioritize within-subject designs that pair median and ulnar stimulation to definitively assess somatotopic specificity. To establish the clinical utility of temporal potentiation, longer-term studies spanning multiple sessions are necessary to monitor habituation over time. Importantly, future investigations should bridge the gap between cortical hemodynamics and functional outcomes by integrating fNIRS with behavioral measures, such as grip force control or perceptual drift [4], [45]. To address individual heterogeneity, adaptive TNS controllers utilizing real-time fNIRS feedback could be explored to modulate stimulation timing and determine whether personalized temporal parameters can stabilize the variable trajectories observed in this study [17], [18].

